# Deterministic approximation for population dynamics in the presence of advantageous mutants

**DOI:** 10.1101/2022.01.04.474956

**Authors:** Ignacio A. Rodriguez-Brenes, Dominik Wodarz, Natalia L. Komarova

**Affiliations:** Department of Mathematics, University of California Irvine, Irvine, California 92697, USA; Department of Population Health and Disease Prevention, University of California, Irvine, California 92617, USA

## Abstract

Spatial stochastic simulations of evolutionary processes are computationally expensive. Here, based on spatially explicit decoupling approximations (SEDA) introduced in [1], we derive a deterministic approximation to a spatial stochastic birth-death process in the presence of two types: the less advantageous resident type and a more advantageous mutant. At the core of this technique are two essential steps: (1) a system of ODEs that approximate spatial interactions among neighboring individuals must be solved; (2) the time-variable has to be rescaled with a factor (called “*α*”) that depends on the kinetic parameters of the wild type and mutant individuals. An explicit formula for *α* is derived, which is a power law of division and death rates of the two types. The method is relatively fast and provides excellent time-series agreement with the stochastic simulation results for the spatial agent-based model. The methodology can be used to describe hard selective sweep events, including the expansion of driver mutations in carcinogenesis, bacterial evolution, and aspects of resistance dynamics.

## 1 Introduction

The processes of spatial colony spread (colonization) and associated evolution are at the core of many biological and biomedical systems, including cancer generation, evolution and spread; the functioning of biofilms; and macroscopic ecological dynamics.

Spatial evolutionary dynamics have received mathematical attention in the recent years, both computationally [2] and theoretically. For example, paper [3] considered a spatially distributed population where mutations were generated and could spread to neighboring locations; time distribution that was needed for an individual to acquire *k* mutations was calculated. Deterministic equations for stochastic spatial games were derived in [4] under the assumption of long-range interactions. In [5], a death-birth process on a lattice was compared with the dynamics of the replicator equation, showing that space significantly modified evolutionary outcomes of the dynamics. These contributions continue a rich tradition of mathematical studies focussing on important statistics of various types of spatial evolutionary stochastic processes, see e.g. [6, 7, 8, 9, 10]; other approaches involve partial differential equation (PDE) modeling, where evolutionary dynamics of traits (e.g. dispersal) are studied in spatial settings [11, 12, 13]. Many important results in this area were obtained in the context of applications to cancer dynamics, see e.g. [14, 15, 16].

Deterministic models are usually not the tool of choice when modeling spatial evolutionary processes. Agent-based modeling is a popular method that allows capturing mutant generation and spread through space. It was used to study, for example, mutant generation in spatially expanding colonies [17, 18], focusing on advantageous [14, 18] and disadvantageous [19] mutants. A number of analytical insights have been generated in the form of scaling laws [20]. These laws for example specify how the number of mutants in a growing system scales with the total number of individuals, given the dimensionality of the system [21]. Another type of results concerns competition dynamics in games, where the selection process is significantly affected by the presence of spatial interactions [22, 23, 24, 25].

When it comes to the description of the actual temporal dynamics of a stochastic agent-based model, analytical approximations are difficult to find. Most PDE methods are based on mean-field behavior and as a consequence they result in inaccurate descriptions of the average trajectories of stochastic agent based models, i.e., they not provide good time series agreement with the agent-based models. In [1] we studied spatial colonization dynamics in the presence of a single species, and derived deterministic approximations of a spatial stochastic process. In the present paper we extend our previous results to a more complex situation where two phenotypes are present, to which we refer as the “wild type” and “mutant” individuals. Here we assume that mutants are advantageous with respect to wild types, and study their spread in the bulk of a “resident”, or “wild type”, population, which undergoes spatial growth and eventually fills out the grid. We develop a deterministic spatially explicit approximation of a two-species 2D stochastic birth-death process implemented as an agent-based model, which provides good time series agreement with the expected behavior of the system. Scenarios that are handled by our methodology are relevant for the theory of hard selective sweeps in population genetics [26], which have been described in cancer [27], bacterial evolution [28], etc.

This paper is organized as follows. In Section 2 we set up a mathematical formulation of the problem and describe spatially explicit decoupling approximations (SEDA), which are a generalization of the approximation proposed in [1] to the case of two species. In section 3 we demonstrate how SEDA can be used successfully in the case of 2D, 2-species dynamics, and discuss how different configurations of the initial conditions could influence the dynamics. Discussion is presented in Section 3.

## 2 Methods

### 2.1 Preliminaries

We begin by considering a birth-death process for two species, *x* and *y*, in a two-dimensional rectangular lattice. In the context of this study one can think of species *x* as wild-type individuals and species *y* as mutants. For any given site in the lattice with coordinates (*i, j*) we introduce the random variables *x*_*ij*_(*t*) and *y*_*ij*_(*t*), which satisfy: *x*_*ij*_(*t*) = 1 and *y*_*ij*_(*t*) = 0 if the site it occupied by an *x* individual at time *t*; *x*_*ij*_(*t*) = 0 and *y*_*ij*_(*t*) = 1 if the site is occupied by a *y* individual; or *x*_*ij*_(*t*) = 0 and *y*_*ij*_(*t*) = 0 if the site is empty (at any given time there can be at most one individual at each site). We consider the *ℓ*^1^ distance in the lattice and by analogy we define the distance between two random variables as the distance between the coordinate sites, for example dist(*x*_*ij*_, *y*_*lk*_) = |*i* − *l*| + |*j* − *k*|. Note that each lattice point has exactly four neighbor sites that lie within a distance of one from it. A site and its four nearest neighbors make up a von Neumann neighborhood of radius 1. We assume that individuals reproduce stochastically onto each unoccupied nearest neighboring site with rates *L*_*x*_*/*4 for *x* individuals and *L*_*y*_*/*4 for *y* types; they also die stochastically at rates *D*_*x*_ and *D*_*y*_.

To simplify the notation we write 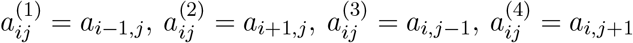, where *a* here stands for an *x* or *y* random variable. If we use angular brackets to denote the expected value, the stochastic process obeys the following equations:

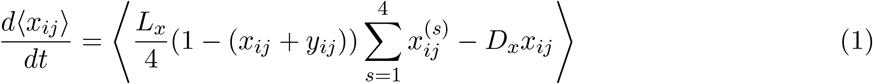

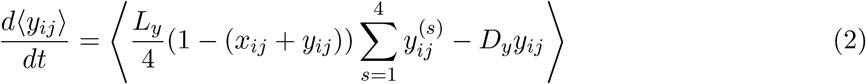

Equations (1) and (2) introduce terms of the form ⟨*ab*⟩, where *a* and *b* are random variables. If we define the functions *X*_*ij*_ = ⟨*x*_*ij*_⟩, *Y*_*ij*_ = ⟨*y*_*ij*_⟩, *A*_*ij*1_ = ⟨*x*_*ij*_*x*_*i*+1,*j*_ ⟩, *A*_*ij*2_ = ⟨*x*_*ij*_*x*_*i,j*+1_ ⟩, *B*_*ij*1_ = ⟨*y*_*ij*_*y*_*i*+1,*j*_ ⟩, *B*_*ij*2_ = ⟨*y*_*ij*_*y*_*i,j*+1_ ⟩*M*_*ij*1_ = ⟨*x*_*ij*_*y*_*i*−1,*j*_ ⟩, *M*_*ij*2_ = ⟨*x*_*ij*_*y*_*i*+1,*j*_⟩, *M*_*ij*3_ = ⟨*x*_*ij*_*y*_*i,j*−1_ ⟩, *M*_*ij*4_ = ⟨*x*_*ij*_*y*_*i,j*+1_⟩, then equations (1) and (2) can be rewritten as:

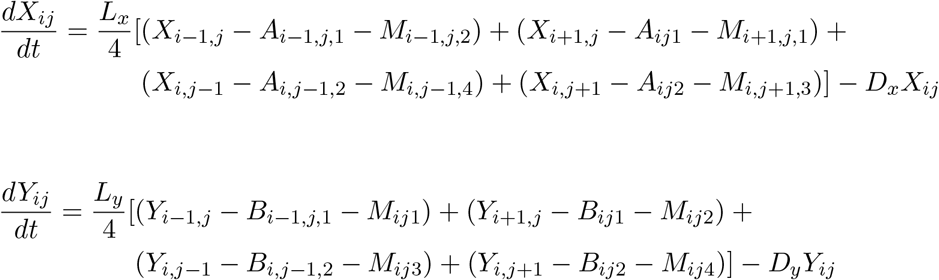

The ODE system defined by the previous equations is not closed. We can expand it by including differential equations for the rate of change of the *A, B*, and *M* variables. These additional differential equations introduce new terms of the form ⟨*bc*⟩ and ⟨*abc*⟩, where *a, b*, and *c* are *x* or *y* random variables that satisfy dist(*a, b*) = dist(*a, c*) = 1, and dist(*b, c*) = 2. In principle, these new terms require additional equations that involve higher order moments. Hence at some point, we need to cut off the process of adding equations and instead use approximations to obtain a closed system.

### 2.2 Spatially explicit decoupling approximations (SEDA)

Here we develop a deterministic spatially explicit approximation for the expected trajectories of the two-species stochastic birth-death process previously described. First, we note that for any triad of random variables, {*a, b, c*}, each of which takes on only the values zero or one, the following relation always holds: ⟨*bc*⟩ − ⟨*abc*⟩ = *P* (*b* = 1, *a* = 0, *c* = 1). If the random variables are defined in the lattice, dist(*a, b*) = dist(*a, c*) = 1, and dist(*b, c*) = 2, we can make the following approximation:

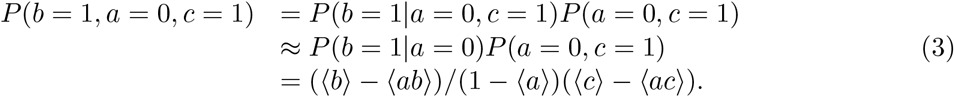

Now, let us look at a specific example on how to use the approximation in (3). The rate of change for *A*_*ij*1_ is:

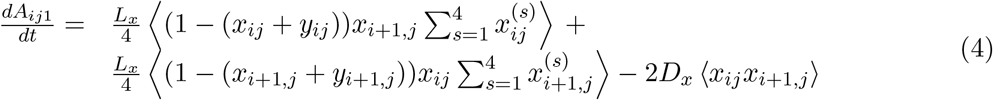

Inside the first pair of angular brackets in equation (4) we find expressions of the form: 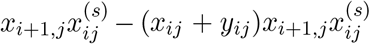. If *s* ≠ 2 and we call *a* = *x*_*ij*_ + *y*_*ij*_, *b* = *x*_*i*+1,*j*_, and 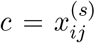, then all three random variables take only the values zero or one, dist(*a, b*) = dist(*a, c*) = 1, and dist(*b, c*) = 2. Hence, the conditions are met to apply approximation (3) to ⟨*bc*⟩ − ⟨*abc*⟩, which yields:

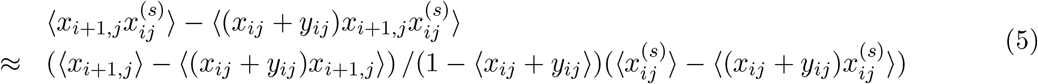

If *s* = 2, then *b* = *c*, and it is easy to see that ⟨*b*^2^⟩ − ⟨*ab*^2^⟩ = ⟨*b*⟩ − ⟨*ab*⟩.

Intuitively, for the spatial stochastic process, the approximation *P* (*b* = 1|*a* = 0, *c* = 1) ≈ *P* (*b* = 1|*a* = 0) embedded in (3) assumes that the probability that a site in the lattice is occupied (*b* = 1) is only weakly dependent on the probability that another site two units of distance apart is also occupied (*c* = 1). However, the fact that individuals arise through replication of nearest neighbors, suggests that on average *P* (*b* = 1|*a* = 0, *c* = 1) *> P* (*b* = 1|*a* = 0). Hence, we can introduce a parameter *ϵ* and set the approximation to:

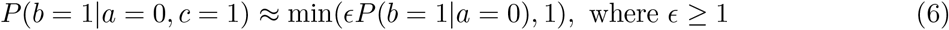

To set the value of *ϵ*, we refer to our previous work related the one species model [REF], where we found a formula for *ϵ* as a function of the ratio *λ* = *D/L*. For the two-species model discussed here we will use the same formula for *ϵ* as a function of *λ*_*y*_ ≡ *D*_*y*_*/L*_*y*_. The reason for choosing *λ*_*y*_ is that under the assumptions of this study, the mutant population is advantageous and will eventually become dominant. More concretely, the explicit formula for *ϵ* used throughout this paper is: 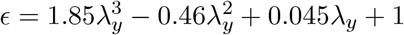.

Finally, we note that the approximation in (3) is only reasonable when ⟨*a*⟩ *<* 1 (otherwise we have *P* (*b* = 1, *a* = 0, *c* = 1) = 0). For this reason, to simplify the notation, we define *δ*_*ij*_ as 1*/*(1 − *X*_*ij*_ − *Y*_*ij*_) if *X*_*ij*_ + *Y*_*ij*_ *<* 1, or zero otherwise. We are then ready to apply he procedure in (5) to the rate of change equations for all the *A, B*, and *M* variables. Doing so we arrive at the following closed system of differential equations, which approximates the expected trajectories of the two-dimensional stochastic birth-death process.

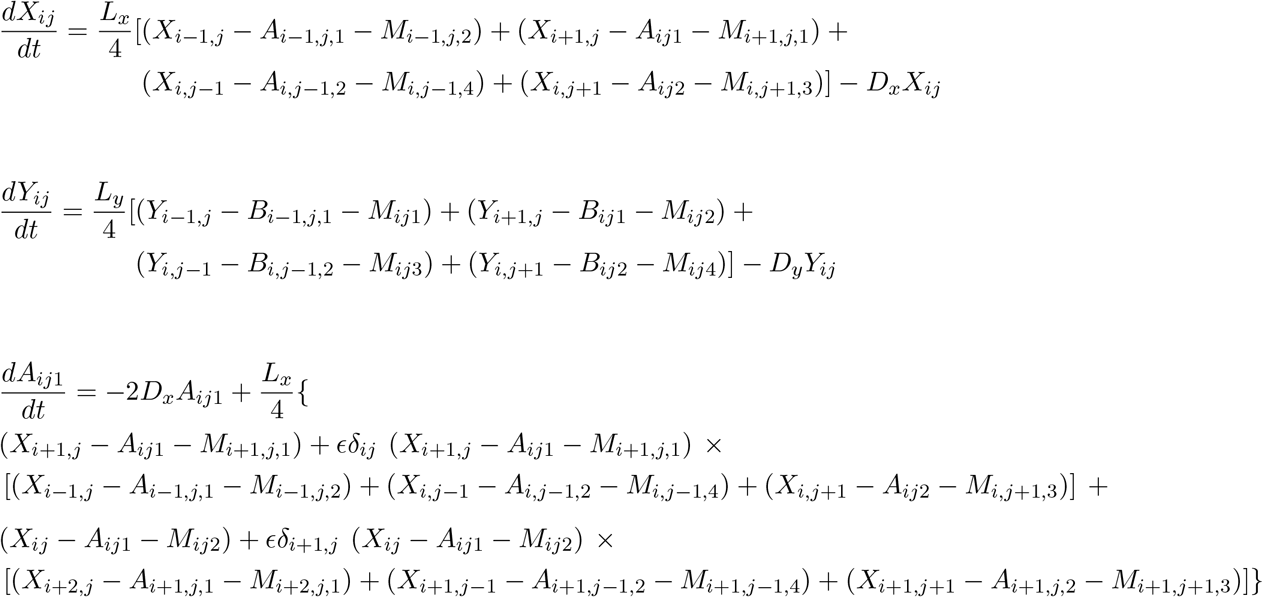

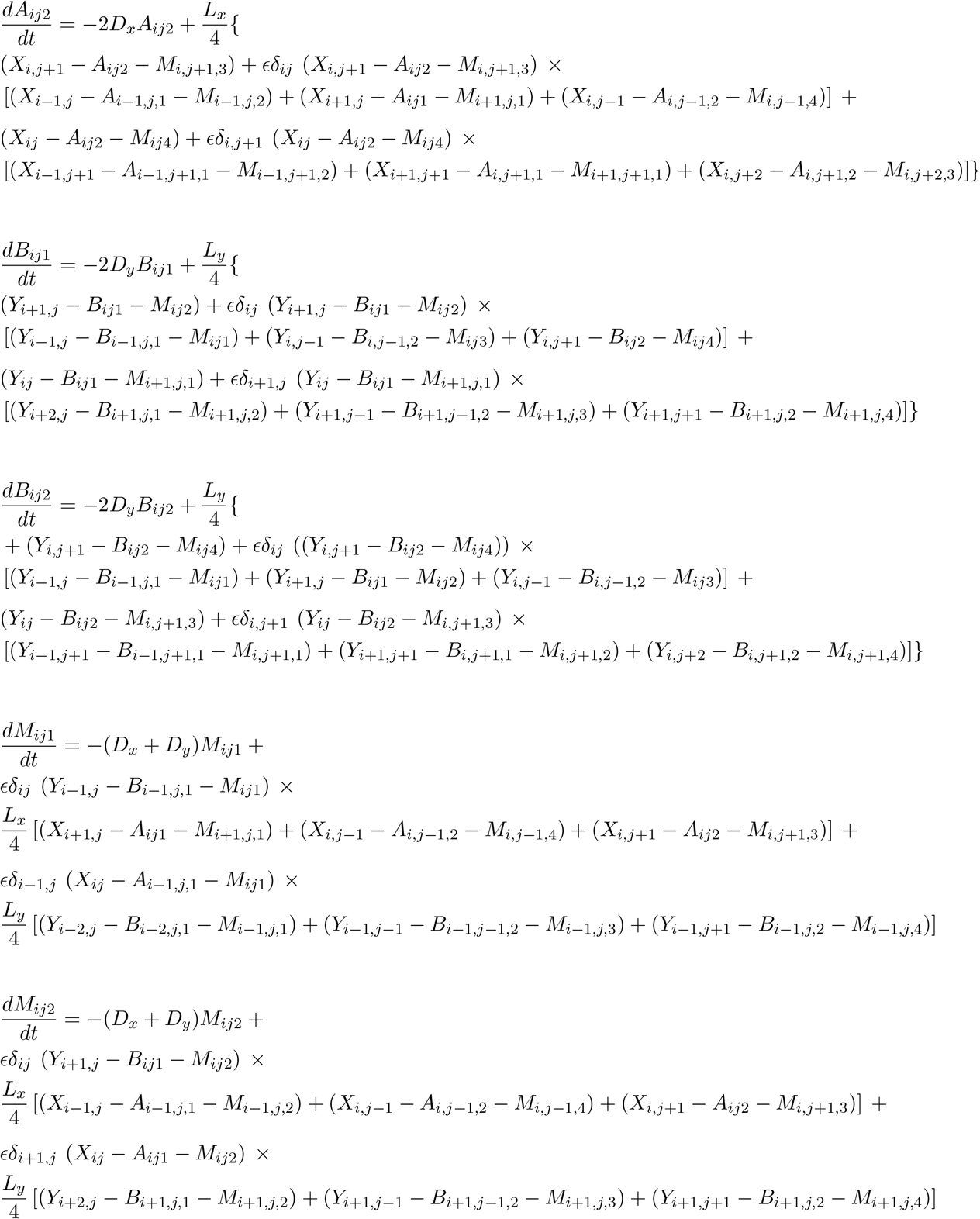

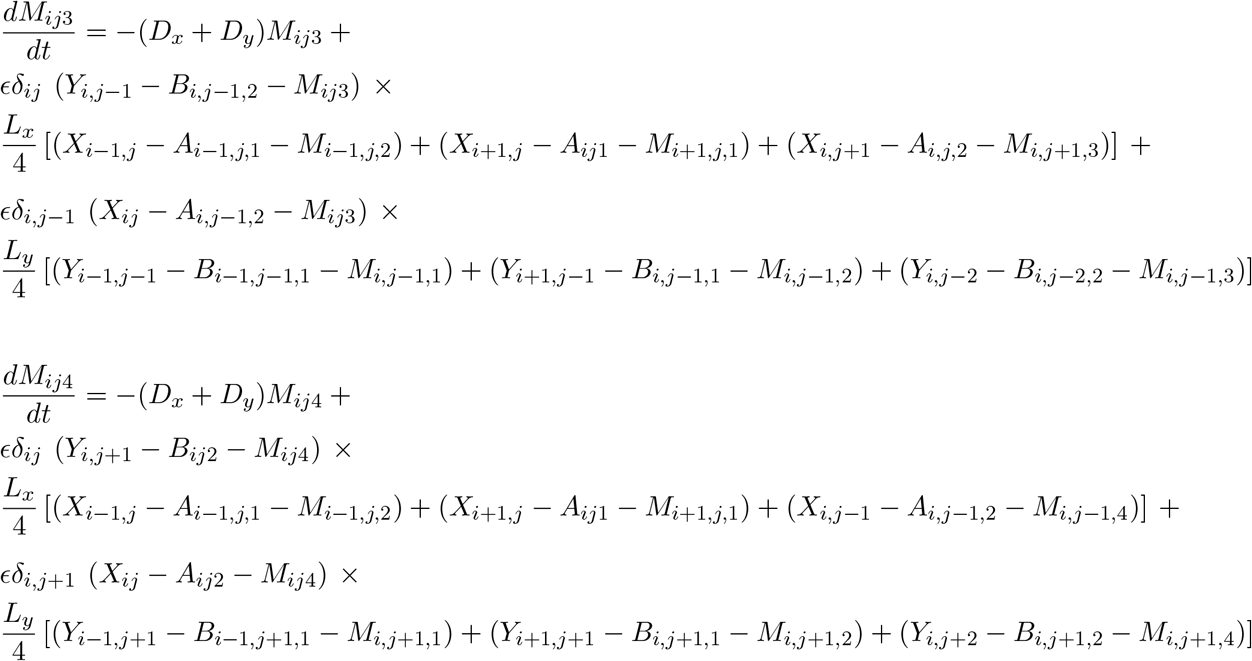

## 3 Results

### 3.1 Time-scaling SEDA and first observations

Before we look at the first set of results, we begin by delimiting the scope of our research and expanding on our methodology. First, we are interested in the application where in an expanding population of homogeneous individuals a second species arises from a mutation. For this reason we limit our discussion to initial conditions where a small number of advantageous mutants, *y*, proliferate amongst a background of a much larger population of wild type individuals, *x*. We define advantageous mutants as those that satisfy *L*_*y*_ ≥ *L*_*x*_ and *D*_*x*_ ≥ *D*_*y*_, with at least one of the previous relations being a strict inequality. Second, we discovered in our work for the one-species model, that a good fit to data from stochastic simulations requires the introduction of a time scaling parameter *α*. This observation remains true for the two-species model. More precisely, if *S* is the total number of individuals and the SEDA ode system yields the functional relation *S* = *F* (*t*), then we can improve the approximation by setting *S*_*α*_ = *F* (*αt*), where the value of *α* is determined through least-squares fitting.

The next step is to look at the possible outcomes for the model, which are exemplified by the panels in Figure 1. Panel (a) shows an instance where there is excellent agreement between the stochastic results and the deterministic approximation. In Panel (b) this agreement is clearly worse. The crucial difference between panels (a) and (b) are the rates (*L*_*x*_, *D*_*x*_, *L*_*y*_, *D*_*y*_). Panel (c) shows two instances with good agreement. Here, the rates are the same, but the location of the initial cluster of mutants differs (located close to the center or edge of the wild type population). The key thing to notice in this panel is that the two values of *α* employed in the plots, *α*_center_ and *α*_edge_, are different. In what follows, we first explore scenarios where initially the mutants are located close to the center of the colony. Later we investigate the role of the initial placement location.

**Figure 1:**
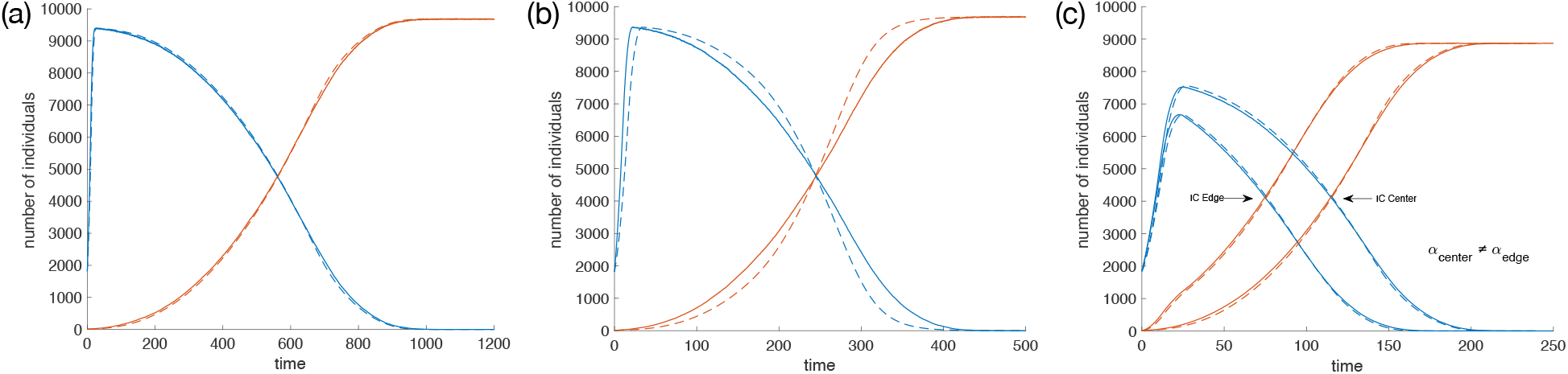
Total number of individuals vs time. Wild type individuals in blue; mutants in red; average from more than 100 stochastic simulation in solid lines; deterministic approximation dashed. (a) *L*_*x*_ = 1, *L*_*y*_ = 0.75, *D*_*x*_ = 0.3, *D*_*y*_ = 0.15, *α* = 1.47. (b) *L*_*x*_ = 1, *L*_*y*_ = 3, *D*_*x*_ = 0.3, *D*_*y*_ = 0.6, *α* = 2. (c) Different values of parameter *α* for different placements of initial cluster of mutants: *L*_*x*_ = *L*_*y*_ = 4, *D*_*x*_ = 0.9, *D*_*y*_ = 0.5, *α*_center_ = 1.48, *α*_edge_ = 1.71.

The previous discussion suggests that we need to establish a domain *D* in the (*L*_*x*_, *D*_*x*_, *L*_*y*_, *D*_*y*_) space where there is good agreement between stochastic and deterministic results. We will do so in the next section, and in the process obtain an explicit formula for *α* that is successful in this domain. Then we will look at the location of the initial cluster of mutants and find that the formula for *α* is valid when the initial mutant cluster is not very close to the edge of the wild type population.

To asses an approximation’s success we introduce a metric that measures the distance between the stochastic results and the approximation. We will also use this metric to verify that enough simulations are carried out to obtain a small confidence interval for the expected trajectories. If (*X*_1_, *Y*_1_) and (*X*_2_, *Y*_2_) are two time series for each species (*x* and *y*), we define the distance *F* between them as follows:

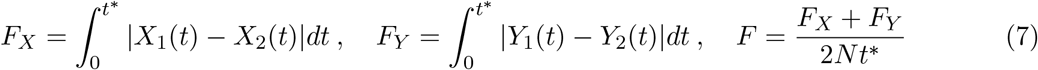

where *N* is the number of sites in the grid, and *t*^*\**^ is the first time when the number of individuals is within 99.9% of the steady state. Note that this metric is based on the *L*^1^ distance between functions, and is normalized by lattice size and the time it takes for the system to reach equilibrium.

### 3.2 Survey of domain *D* and formula for *α*

We are looking for a domain (or set of rates) *D* where the deterministic method accurately approximates the average behavior of the stochastic process. First, from our previous work on the one species model with rates *L* and *D*, we found that the approximation was very successful when *D/L* ≤ 0.4, which roughly corresponds to steady-state densities ≥ 50%. Hence, if we call 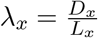 and 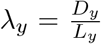, we limit our domain search to rates that satisfy *λ*_*x*_, *λ*_*y*_ ≤ 0.4. Note also that since time units in the model are arbitrary, we can assume without loss of generality that *L*_*x*_ = 1. With this assumption we then limit the range of *L*_*y*_ to: 1 ≤ *L*_*y*_ ≤ 1.2. This is equivalent to a maximum of a 20% advantage in the division rates for mutants, which we argue is not very restrictive, since most advantageous mutations probably provide only modest improvements in the kinetic rates. The relations *L*_*x*_ = 1, 1 ≤ *L*_*y*_ ≤ 1.2, *D*_*x*_ ≤ *D*_*y*_, and *λ*_*x*_, *λ*_*y*_ ≤ 0.4 fully determine our proposed domain *D*.

As a next step we asses the quality of the approximation in *D*. To accomplish this we discretize the domain by considering all triplets (*L*_*y*_, *D*_*x*_, *D*_*y*_) where: (i) *L*_*y*_ takes values in {1, 1.05, 1.1, 1.15, 1.2}; (ii) *D*_*x*_ and *D*_*y*_ take values in {0, 0.05, 0.1, 0.15, 0.2, 0.25, 0.3, 0.35, 0.4}; and (iii) *D*_*y*_ ≤ *D*_*x*_ and *L*_*x*_ ≤ *L*_*y*_ (where at least one inequality is strict). This results in 212 points (rate combinations).

To generate the initial conditions for the simulations we do the following: First, for each of the discrete values of *D*_*x*_ we place a single wild type individual at the grid’s center and let the population grow until it forms a random spatial cluster with at least 1800 individuals. Then for each point in *D* we select as initial conditions the random cluster corresponding to the wild type death rate, and a small cluster of 29 mutants arranged as a disk at the grid’s center. Finally, to estimate the average trajectories from the stochastic process we perform for each point in *D* enough simulations to obtain a distance *F* ≤ 0.03, between the curves for the upper and lower bounds of the 95% confidence interval. This threshold, *F* = 0.03, results in a very small 95% confidence interval at each time point for the expected number of individuals (Figures 2c and 2d). Once we have the expected trajectories from the stochastic process, we use the ODEs with a scaling factor, *α*, to find the best fits to the mean stochastic trajectories. This results in 212 values of *α* that correspond to different sets of kinetic rates. We find that the deterministic approximations provide excellent agreement with stochastic results. Indeed, for all 212 points tested we found a distance between stochastic and deterministic results *F <* 0.032. Figures 2a and 2b depicts the tested points with the highest and lowest agreement based on the metric *F*.

**Figure 2:**
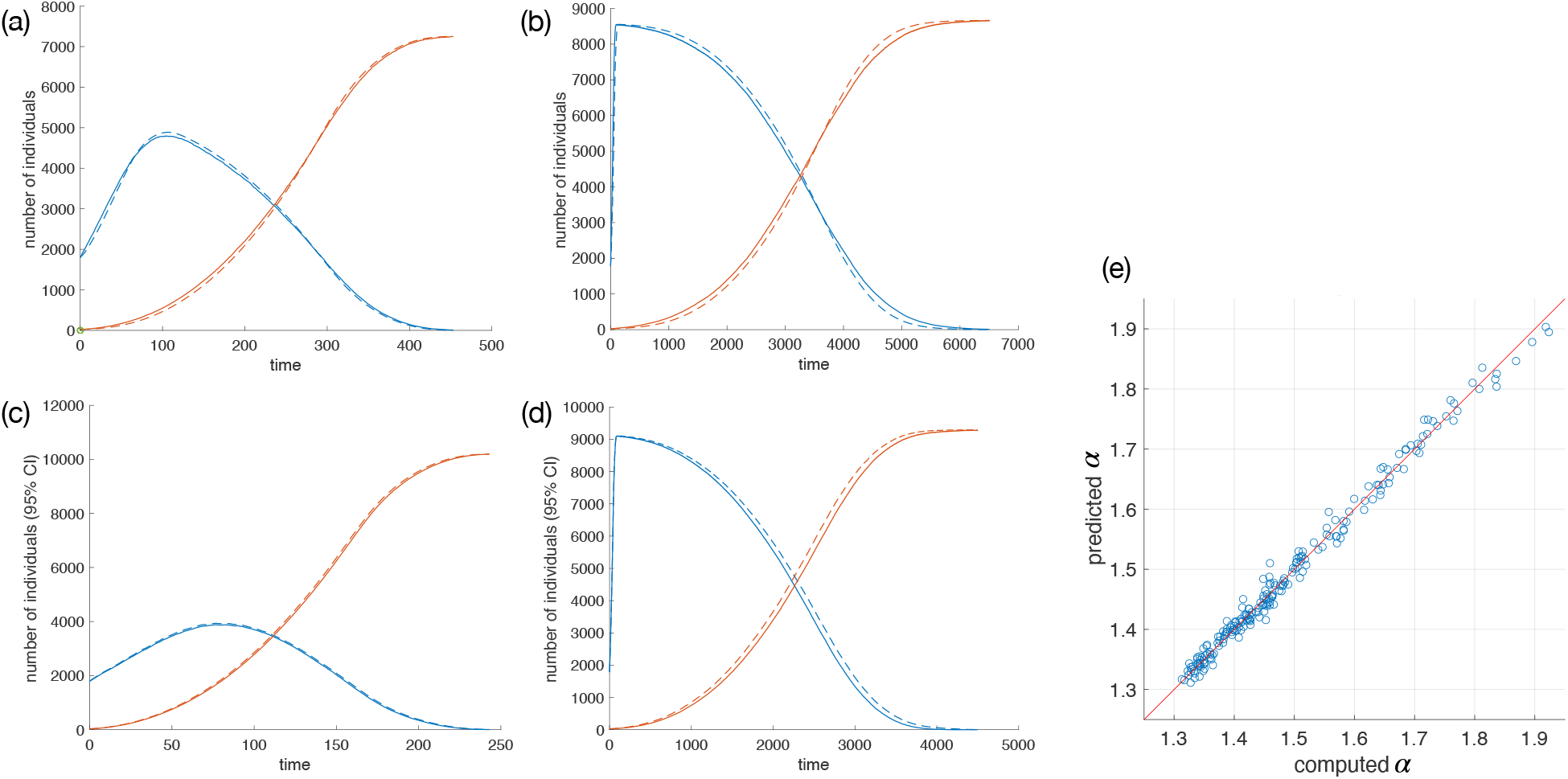
(a) and (b), best and worst fits for the 212 parameters in domain *D*, according to the metric *F*. Wild type in blue; mutants in red. Stochastic average in solid lines; deterministic approximation dashed. (a) Best: *D*_*x*_ = 0.4, *L*_*y*_ = 1.15, *D*_*y*_ = 0.3, *F* = 0.013. (b) Worst: *D*_*x*_ = 0.05, *L*_*y*_ = 1.05, *D*_*y*_ = 0.05, *F* = 0.03. (c) and (d), widest and narrowest 95% confidence intervals for the stochastic simulations in *D*. Upper bound dashed; lower bound solid. (c) Narrowest confidence interval: *D*_*x*_ = 0.4, *L*_*y*_ = 1.05, *D*_*y*_ = 0, *F* = 0.010. (d) Widest: *D*_*x*_ = 0.1, *L*_*y*_ = 1.15, *D*_*y*_ = 0.1, *F* = 0.032. In all panels *L*_*x*_ = 1. (e) Computed values of *α* vs. the values predicted by the linear regression formula. Each circle corresponds to a different parameter set in domain *D*.

We would now like to derive an explicit formula for *α* in terms of the kinetic rates *L*_*y*_, *D*_*x*_, and *D*_*y*_. For this we propose a polynomial function,

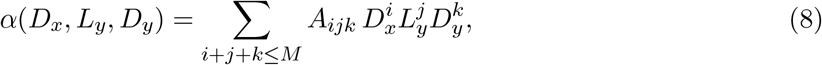

where the coefficients *A*_*ijk*_ are constants and *M* is the polynomial degree. This formula is linear in the coefficients, so we can apply linear regression to the data obtained from the discretization of D. We call the resulting model the full regression model of degree *M*. Next, we apply ANOVA to the full regression model; discard the terms with *p*–values greater than 0.05; and rerun the regression without these terms. We continue this process iteratively until we are left with a model with only significant terms, which we call the reduced regression model of degree *M*. Finally for model selection, we apply Akaike information criterion with small sample size correction (AICc) to the full and reduced regression models with degrees ranging from three to six. Using Akaike’s weights we find that compared with the other models, the 5th degree reduced model minimizes information loss with a normalized probability of 0.97. This model yields a polynomial formula for *α* with 33 terms specified in Table 1. The formula provides remarkable agreement with the computed values for *α*, as shown in Figure 2e. For more details, please see Appendix A.

**Table 1:**
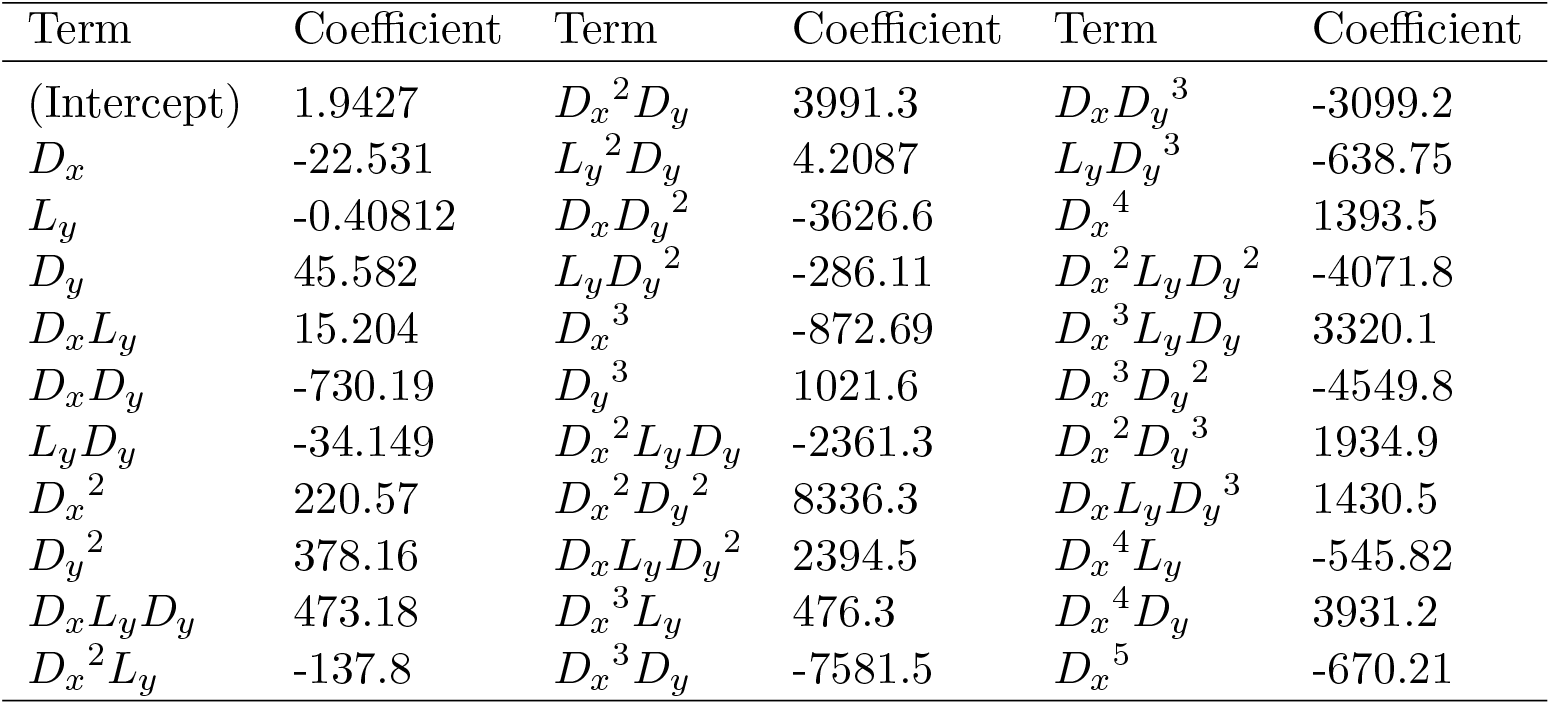
Formula for the 5th degree reduced model 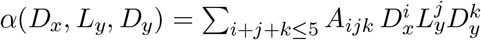 non-zero coefficients shown.

### 3.3 Validity of the formula for *α* away from the boundary

As shown in figure 1c, the values of *α* that provide the best fitting curve can be different depending on the initial location of the mutant cluster. In particular, its proximity to the boundary of the wild type population seems to play a role. To investigate this phenomenon, we first formally define a population’s boundary and then study the applicability of formula (8) with coefficients given in Table 1, depending on the distance of the initial mutant cluster to this boundary.

In the model there is a simple way to characterize the boundary of a single species population. Let *G* be the set of occupied sites in the lattice. If the lattice is large enough we can select an empty site *e* that is clearly outside the population mass; then, we can define the outside set, *O*_*G*_, as the largest connected set of unoccupied sites that contains *e*; as usual a connected set is one in which any two elements in the set can be connected by a path that lies entirely within it. We could then define the boundary of *G* as the set *B*_*G*_ made up of all elements of *O*_*G*_ that have a nearest neighbor in *G*. We call *B*_*G*_ the type I boundary of *G*.

The problem with type I boundaries is that the when the population density is small, the border set can, through a narrow gap, extend deep into the population, close to its center of mass (Figure 3). However, we can verify that for initial mutant clusters located near the center of the wild type population, the estimates for *α* derived in the previous section are valid, regardless of their distance to the type I boundary. Therefore, we need a boundary definition that provides a more convex envelop of the wild type population. We can define such a boundary algorithmically. First, let 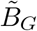 be made up of all the sites in *B*_*G*_ plus all empty sites that have all nearest neighbors in either *G* or *B*_*G*_. Next, we define the set *G*1, which is made up of all the members in *G*, plus all sites in 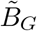 that have two or more nearest neighbors in *G*, or all neighbors in 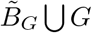. We can then consider the type I border of *G*1, *B*_*G*1_ (note that *G* ⊂ *G*1 guarantees that *B*_*G*1_ envelops *G*). The previous steps are aimed at closing very narrow gaps in the outer regions of the population. Also, by adding to *G*1 points of 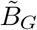 that have two or more neighbors in *G*, the algorithm is working towards a more convex shape of the final boundary. Finally, we can introduce the type II boundary of *G, BB*_*G*_, which is made up of all elements in *B*_*G*1_ that have at least one nearest neighbor in the outside set 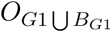. This final step removes unnecessary elements in *B*_*G*1_ that might have been introduced during the previous steps. Figure 3 shows a random cluster with the two types of boundaries for a low density population. In Appendix B we discuss how the type II boundary is better suited for our purposes and from here on we refer to it as simply the boundary.

**Figure 3:**
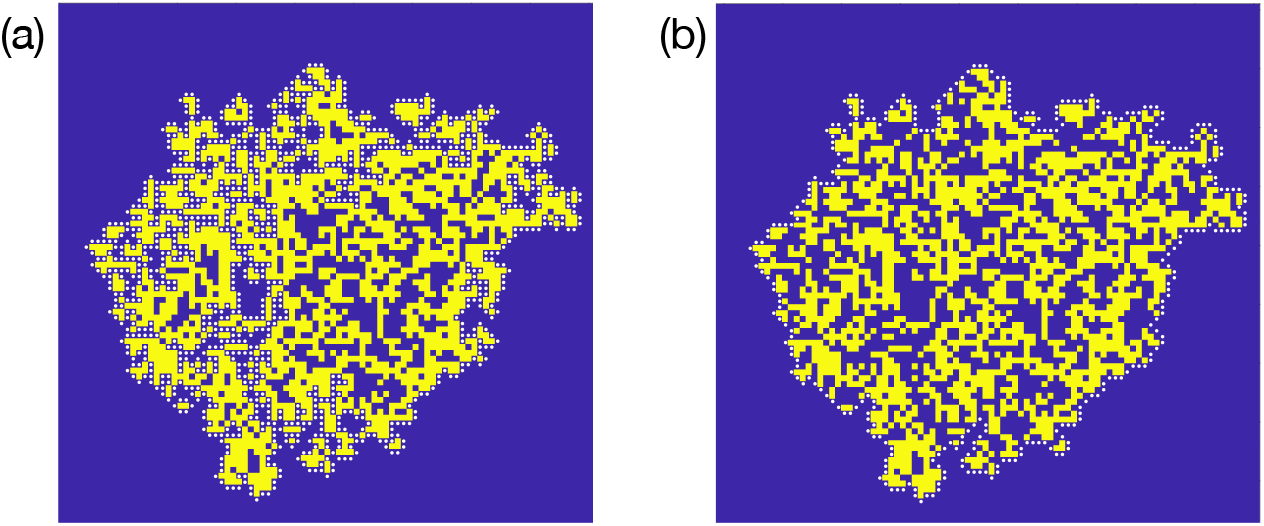
Types I and II boundaries for a random cluster. Occupied sites yellow; empty space in blue; boundary sites with white dots. (a) Type I boundary. (b) Type II boundary. Parameters: *L*_*x*_ = 1, *D*_*x*_ = 0.4.

We are now ready to explore how the location of the initial mutant cluster affects the scaling *α*. To do so, we begin with the same wild type random clusters used as initial conditions in the previous section, and measure the distance between each occupied site and the population’s boundary. For each recorded distance, we then randomly select four individuals that lie at this distance and place in their locations the center of the initial mutant clusters, which once again consist of 29 individuals arranged as a disk. We then calculate the optimal *α* for each of these initial conditions using the methodology described in the previous section. Figure 4 plots simulation results corresponding to high and low wild type population densities (*D*_*x*_ = 0.15 and *D* = 0.35). Here, the horizontal axes specify the distances from the center of the initial mutant clusters to the wild type population’s boundary. We first note that very close to the boundary, the distance between the upper and lower bounds of the 95% confidence interval, *F*, can be larger than our specified threshold *F* ≤ 0.03 (shaded in light blue, panels (a) and (d)). This suggests that when the initial mutant cluster is located very close to the edge, there can be great variability among the stochastic trajectories. Close to the edge we can also observe significant disagreement between deterministic and stochastic results (shaded outcomes in panels (b) and (e)). However, when the initial mutant cluster is located sufficiently far from the edge (more than 9 or 11 units away), the deterministic approximation provides a good fit. We also note that when the initial mutants are far enough from the boundary, the computed optimal *α*’s for these simulations are remarkably close to those found in the previous section, referred to as *α*_0_ in panels (c) and (f) (a value of *α/α*_0_ = 1 indicates exact agreement). In all the parameters we tested, we found that when the initial cluster of mutants is located at least 11 units of distance away from the boundary, the *α*’s found in the previous section are valid, and the corresponding deterministic approximations provide a very good agreement with stochastic results.

**Figure 4:**
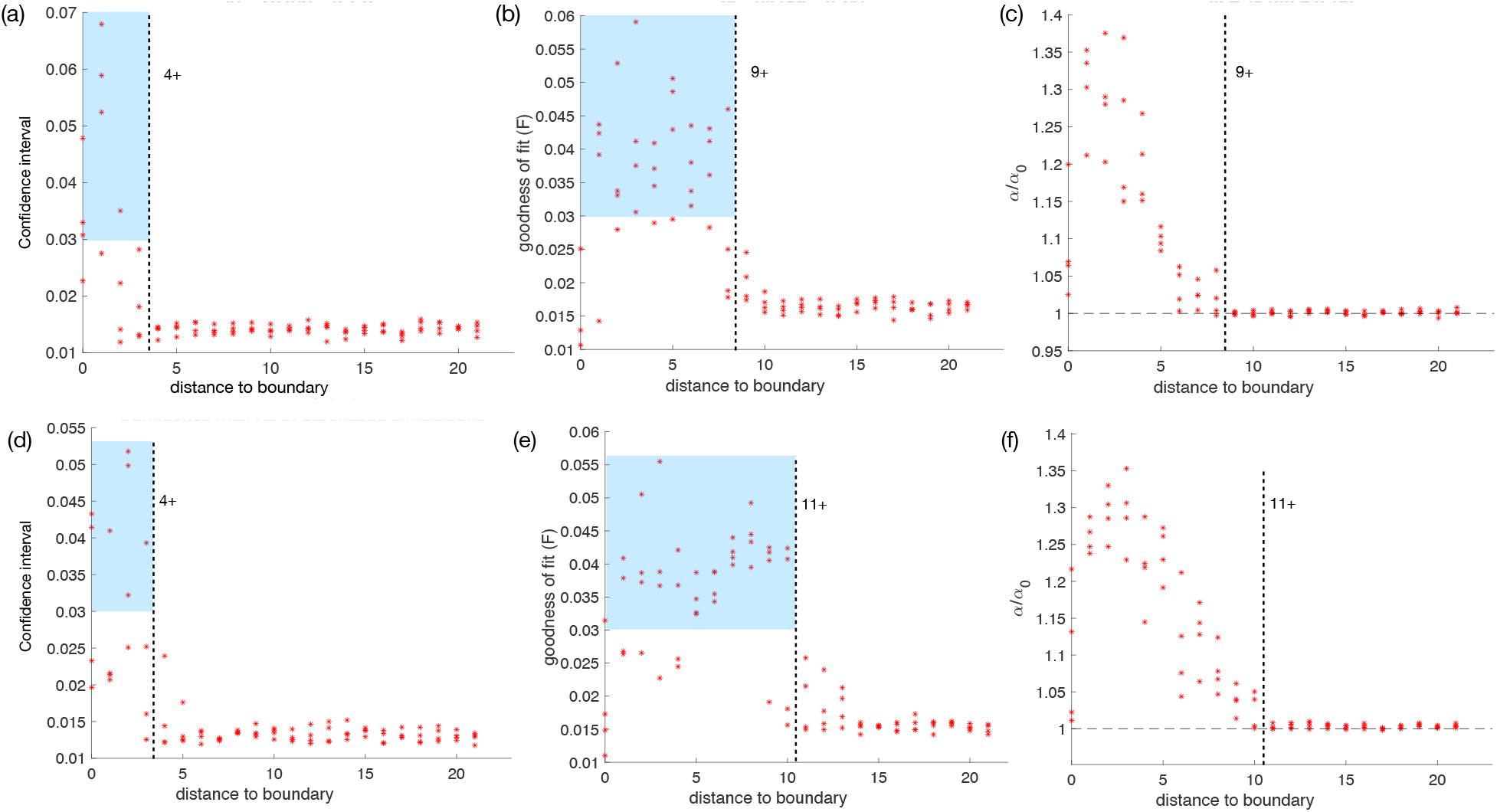
Effects of distance from the initial mutant cluster to the boundary. First row: *D*_*x*_ = 0.15, *L*_*y*_ = 1.05, *D*_*y*_ = 0.05. Second row: *D*_*x*_ = 0.35, *L*_*y*_ = 1.05, *D*_*y*_ = 0. Each point is the result of 100 simulations. (a) and (d) Distance between the upper and lower bounds of the 95% confidence interval. (b) and (e) Goodness of fit between stochastic results and the approximation. Shaded region indicates the areas where the distance *F >* 0.03 (our threshold). (c) and (f) Ratio between the optimal *α* for each specific location of the initial conditions, and the alphas calculated during the survey of domain *D*. See text for discussion.

### 3.4 Application to large population sizes

We are interested in testing our model in much larger grids and with larger clusters as initial conditions. If the initial clusters are large enough, then the stochastic trajectories of the total number of individuals as a function time behave almost deterministically; and hence, a single stochastic run is sufficient to track the time evolution of the expected number of individuals. To generate this type of initial conditions we start with a single wild type individual and using an agent-based model let the population grow into a random cluster. At some point when the population ≥ 20000 we randomly choose an individual for mutation (located at least 11 units of distance away from the boundary). The simulation then continues until the mutant population becomes “large” (here defined as 10,000 or more individuals); the state of the system at this point is the large cluster to be used as initial conditions. Having a large cluster as the initial conditions, we then compare the output of a single stochastic simulation with the results from the deterministic approximation that uses the formula for *α*. Figure 5a shows a set of initial conditions generated through this procedure. In this figure: wild type individuals are red, mutants are cells are green, and empty space is black. Both cell types have the same birth rates, but different death rates. Figures 5b and 5c depict snapshots of the system at a same future time *t*_1_, generated by a fully stochastic simulation (Figure 5b) and the deterministic approximation (Figure 5c). Figure 6 shows the time series for the total number of individuals for a different set of parameters calculated again in a large grid with large clusters as initial conditions. It shows excellent agreement between results from a stochastic agent-based simulation and the deterministic approximation.

**Figure 5:**
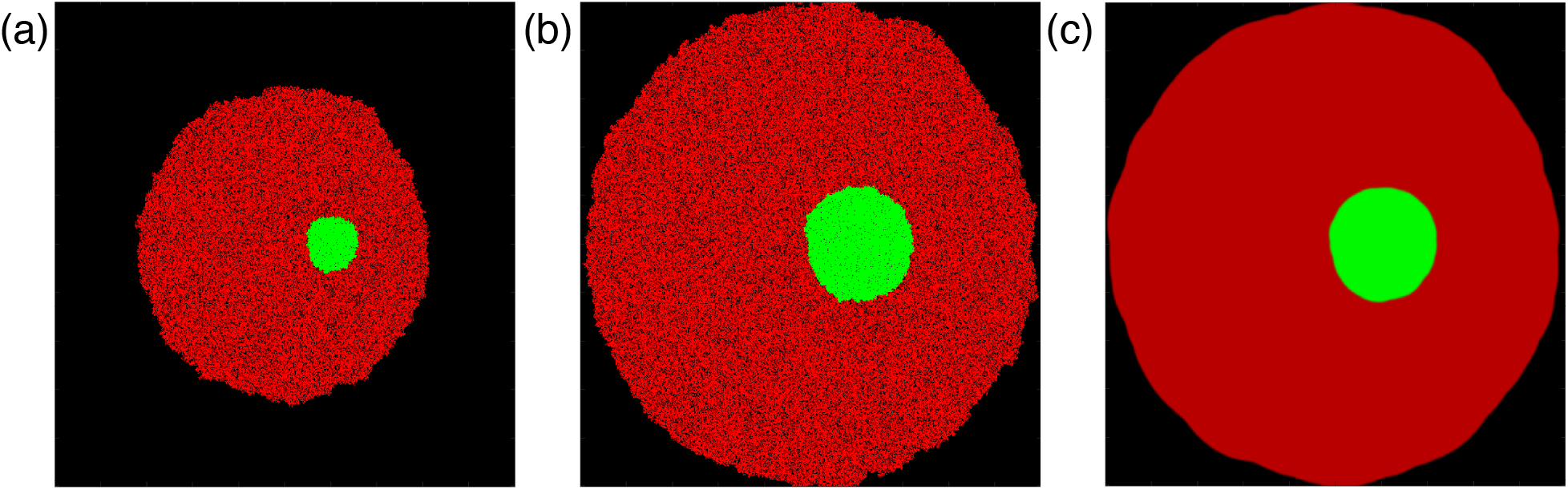
2D results from a stochastic simulation and the deterministic approximation. (a) Initial conditions: wild type red; mutants green; and empty space black. (b) State of the system at a future time, *t*_1_, generated by an agent-based simulation. (c) State of the system at the same time, *t*_1_, computed by solving the deterministic approximation. *L*_*x*_ = *L*_*y*_ = 1, *D*_*x*_ = 0.25, *D*_*y*_ = 0.025.

**Figure 6:**
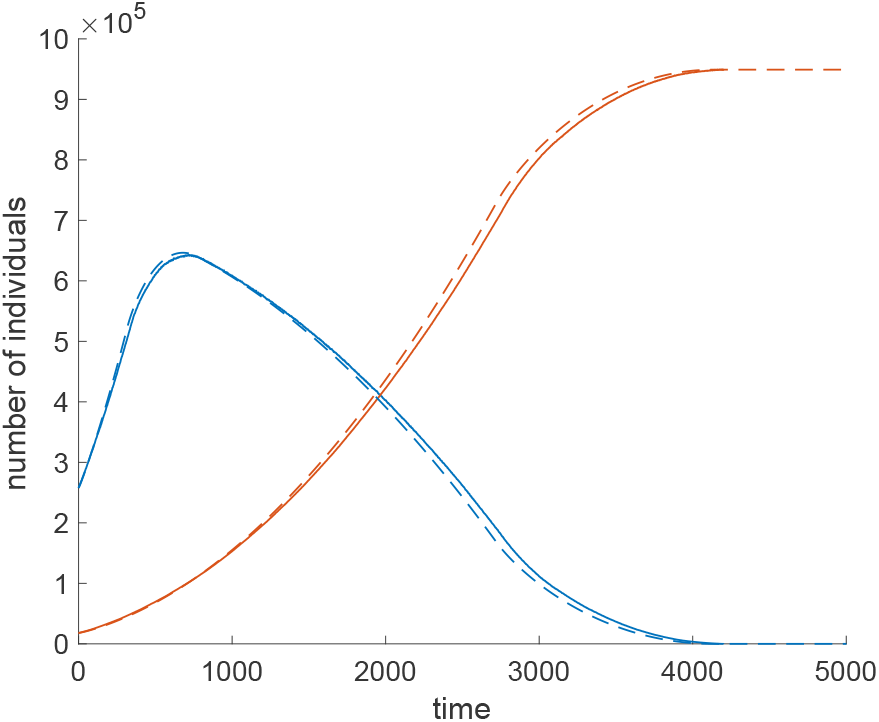
Time series results for a large grid. A small mutant cluster is placed near the center of larger random cluster of 25000 wild-type individuals. Stochastic simulations proceed until the mutant population reaches 18000, when this happens the state of the system is recorded and used as the initial conditions. The picture depicts results from a single stochastic simulation (solid lines) and the deterministic approximation (dashed). Grid size: 1000 × 1000. Parameters: *L*_*x*_ = *L*_*y*_ = 1, *D*_*x*_ = 0.25, and *D*_*y*_ = 0.05.

Finally, we discuss the numerical solution of the approximate systems. Consistent with our findings for the one species model [1], we found that a forward Euler step size of *dt* = 0.0125*L*_*m*_, where *L*_*m*_ = max(*L*_*x*_, *L*_*y*_), yields good results in domain *D* for 0.25 ≤ *λ*_*x*_, *λ*_*y*_ ≤ 0.4, and a step size of *dt* = 0.025*L*_*m*_ is sufficient for all *λ*_*x*_, *λ*_*y*_ ≤ 0.4 in the same domain. We observed that at a 10^7^ grid size, the deterministic method (under the forward Euler method) is 2.5 faster, compared to (highly optimized) stochastic simulations; we expect that using a higher order implicit methods, such as Runge-Kutta, for solving the ODEs, would result in an even faster performance of our deterministic method.

## 4 Discussion

In this paper we have developed a technique that allows for an accurate description of spatial dynamics of expanding colonies that consist of two types of individuals, the wild types and advantageous mutants. This is a generalization of our earlier work (see [1]) where we introduced spatially explicit decoupling approximations (SEDA) in the context of one-species colonization dynamics. This deterministic methodology allows for relatively fast computations compared to stochastic agent-based modeling (ABM) simulations that require many repeats to obtain the expected behavior.

This technique contains two essential steps. First, one needs to solve a system of ODEs that approximate spatial interactions among neighboring individuals must be solved; this system is derived here for the von Neumann 2D grid; this can be adapted to other grids using similar methods. Then, the time-variable has to be rescaled with a factor (which we called “*α*”) that depends on the kinetic parameters of the wild type and mutant individuals. An explicit formula for *α* was derived, which is a power law of division and death rates of the two types. The method provides excellent agreement with the (notoriously expensive) stochastic simulation results for the spatial ABM.

The advantage of this method is that it gives accurate predictions not only for the asymptotic behavior (i.e. the correct steady state levels of the mutants after they take over and the whole grid is colonized) but also for the time-series of the colony sizes, both for advantageous mutants and for the wild types that undergo the process of spatial expansion. In this scenario, the mutants that are initially in a small minority, grow inside the bulk of the host species. This is a process that is important for example in the context of selective sweeps [29] experienced by tumors that acquire a new driver mutation. Drivers have been defined as those genetic (or epigenetic) modifications that are essential for the process carcinogenesis and confer selective advantage to cells [30, 31, 32]. The model is directly applicable to such situations and can for example provide information on the timing of the growth of mutant colonies.

The setting of our model can be also be applied to some aspects of the co-dynamics of susceptible and resistant mutants in the context of cancer drug treatments. For example, it has been suggested that mutations that render chronic lymphocytic leukemia (CLL) cells resistant to the drug Ibrutinib could have selective advantage even in the absence of treatment [33]. In such a scenario, our methodology can be used to estimate the timing of resistant mutant growth before the start of treatment. In the opposite case when resistant mutants are disadvantageous in the absence of treatment, our method applies to the after-treatment dynamics, when the resistant cells become the majority, and susceptible cells (that are advantageous compared to the mutants) start growing from relatively small numbers, and in the absence of treatment will eventually take over. Kinetic parameters of cancer cells may be vastly different in different patients [34], and measuring those parameters is a step toward personalized medicine where treatment outcomes are predicted based on patient-specific values. In this context, our methodology can provide a tool for generating expected trajectories for tumor cell dynamics in individual patients.

Our methodology can only be viewed as an initial step toward this overarching goal. At present, it has only be developed for two-dimensional settings. While expanding the methodology to 3D grids is not conceptually difficult, it will lead to an increased complexity of the resulting ODE system. Furthermore, our method gives reliable predictions for most initial locations of the mutant cells, but it is not applicable in the case where the initial mutant colony is at the very front of the expanding host population. A generalization of this method to such scenarios is subject of future work. Finally, a disadvantage of this model is its complexity in terms of the ODE derivation. In this paper the equations were derived for a birth-death process of two coexisting cell types. Increasing the number of types would result in a significant increase of complexity. For example, in principle our approach can be generalized for hierarchical populations of cells that consist, say, of stem and differentiated cells, where some given rules of cell fate decisions are assumed. This however cannot be read off in a straightforward way from the equations used here, but has to be derived by using similar methods.

To conclude, we provided a methodology that allows prediction of the expected trajectories for advantageous mutants that arise in the bulk of the expanding resident population. This contributes to our ability to quantify the dynamics of selective sweeps in different biological contexts.

## Acknowledgements

The authors greatfully acknowledge partial support from the NSF/DMS grant 1662146 and from The NSF-Simons Center for Multiscale Cell Fate Research.

## A Model selection procedure for the polynomial function *α*

Akaike information criterion with small sample correction is given by eq. (9), where 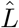 is the maximum value of the likelihood function for the model, *n* denotes sample size, and *k* is the number of parameters. The small sample correction should be used when *n/k* ≤ 40 [REF].

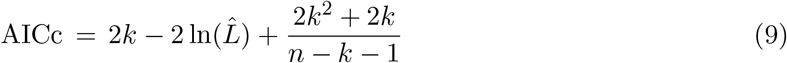

There is a simple way to calculate 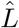. First, we assume that residuals are normally distributed with mean zero, which is strongly suggested by the data (see for example Figure X). Then for linear regression models with *N* (0, *σ*^2^) distributed residuals, 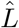 is equal to the the value of the normal likelihood function with mean zero and variance *σ*^2^ evaluated at the residuals from the ordinary least square solution.

The Akaike weights are defined as usual. Let AICc_*i*_ be the value corresponding to model *i*, and AICc_min_ the minimum Akaike score. Then if Δ_*i*_ = AICc_*i*_−AICc_min_, the Akaike weight for model *i* is 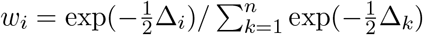, where *n* is the number of models considered. The normalized probability that model *i* model minimizes information loss compared to the other models is then 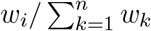 [REF].

## B Domain boundary definition: additional considerations

For some populations it is common to find individuals that are at a very short distance from the type I boundary, but much farther away from the type II boundary. Given that the validity of the formula for *α* depends on the distance from the boundary, the question arises: Is the formula for *α* valid for mutant clusters which are very close to the type I boundary, but farther away from the type II boundary? If the answer is yes, then clearly type II boundaries are preferred for our purposes.

To research the previous question we generated a random wild type population starting from a single individual at the grid’s center with birth rate *L*_*x*_ = 1 and death rate *D*_*x*_ = 0.4. This population is shown in Figure 3. Heat maps of the distance from each occupied site to the type I and type II boundaries are shown in Figures 8(a) and 8(b). We then considered the 110 individuals that had the greatest difference between their distances from the type I and type II boundary. We then proceeded to place clusters of 29 mutants centered at these locations and select from them those clusters that where at most 2 units apart from the type I boundary (these same clusters where at least 17 units apart from the type II boundary). This procedure resulted in 96 mutant clusters whose centers are specified in yellow in Figure 8(c) along with the the type I boundary (white dots). For each cluster we then performed 100 simulations and calculated the optimal alphas for mutants with parameters *L*_*y*_ = 1 and *D*_*y*_ = 0.05. Finally, we compared these optimal alphas with the prediction *α*_0_ = *α*(0.4, 1, 0.05) specified by the formula. We found that every optimal alpha from these simulations deviated from *α*_0_ by less than 0.6% (an excellent agreement). This example thus shows that many mutant origin locations that would be excluded from the validity of the *α*-formula if the the type I boundary is used, are however valid, and are as such classified under the boundary II definition. The behavior just described is typical of what we have found in our simulations, and therefor conclude that the type II boundary is better suited for the purposes and findings of this study.

**Figure 7:**
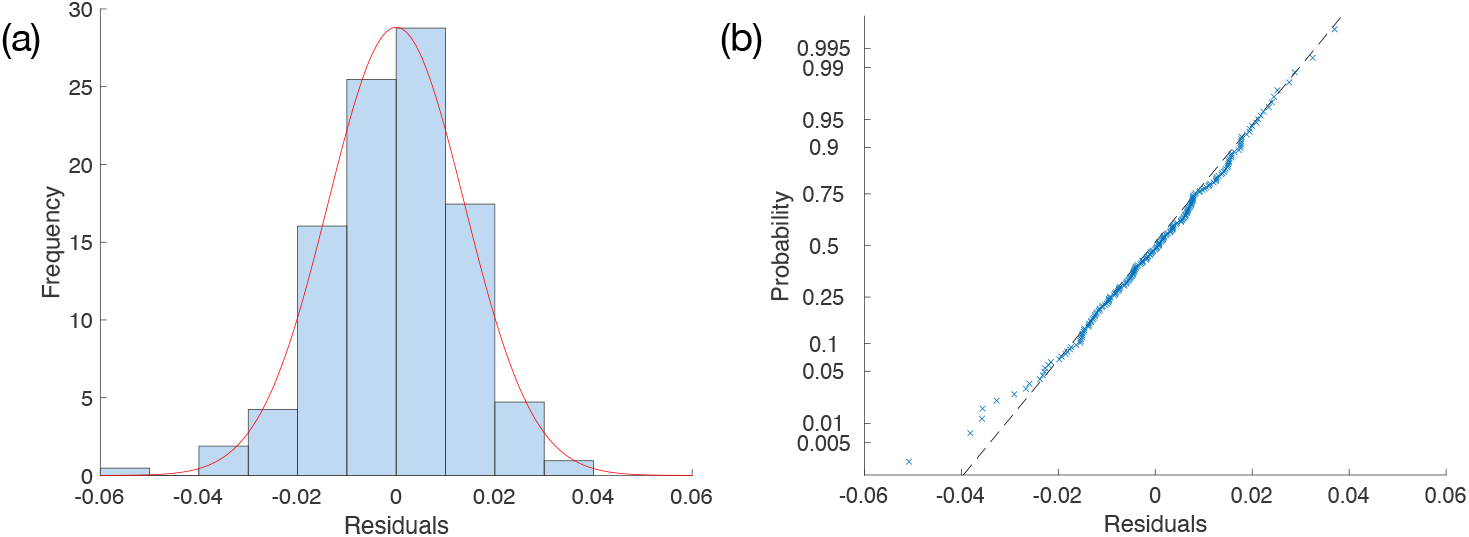
(a) Histogram of residuals from the 5th degree reduced model (bars), compared to plot of normal distribution with same variance as residuals and zero mean (red line). (b) Normal probability plot of residuals for the same model.

**Figure 8:**
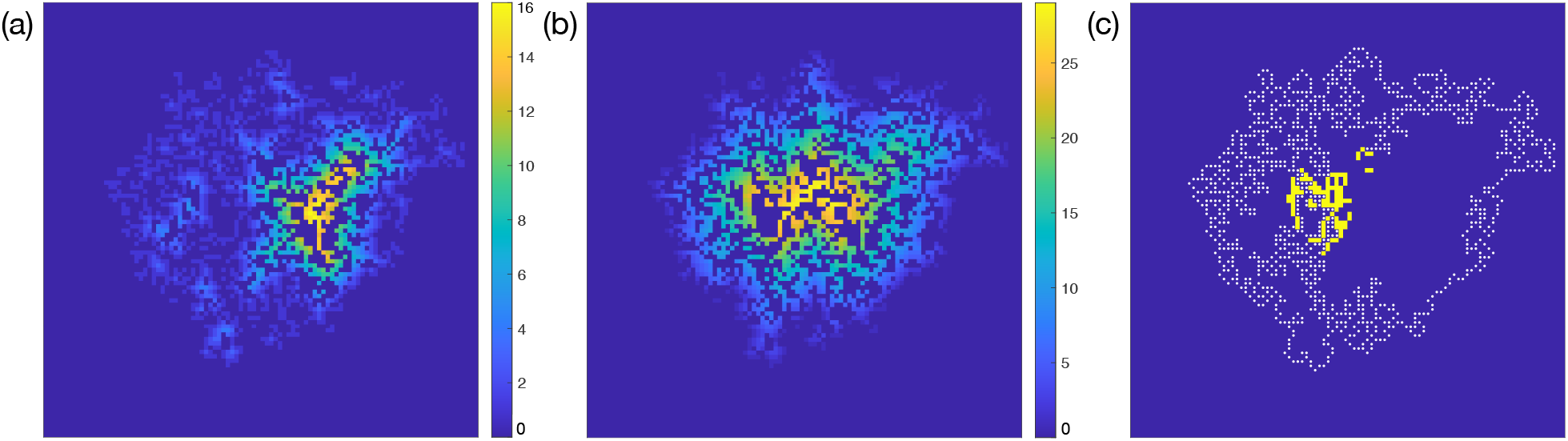
(a) Heat map of distance from occupied sites to the type I boundary. (b) Heat map of distance from occupied sites to the type II boundary. (c) Type I boundary (white dots); in yellow, locations of centers of mutant clusters tested –see text for discussion.

## References

[1] Ignacio A Rodriguez-Brenes, Dominik Wodarz, and Natalia L Komarova. Beyond the pair approximation: Modeling colonization population dynamics. Physical Review E, 101(3):032404, 2020.

[2] Marco Tomassini. Spatially structured evolutionary algorithms: Artificial evolution in space and time. Springer, 2006.

[3] Jasmine Foo, Kevin Leder, and Jason Schweinsberg. Mutation timing in a spatial model of evolution. Stochastic Processes and their Applications, 130(10):6388–6413, 2020.

[4] Sung-Ha Hwang, Markos Katsoulakis, and Luc Rey-Bellet. Deterministic equations for stochastic spatial evolutionary games. Theoretical Economics, 8(3):829–874, 2013.

[5] Stephen Evilsizor and Nicolas Lanchier. Evolutionary games on the lattice: death-birth updating process. Electronic Journal of Probability, 21:1–29, 2016.

[6] Nicolas Champagnat and Sylvie Méléard. Invasion and adaptive evolution for individual-based spatially structured populations. Journal of Mathematical Biology, 55(2):147–188, 2007.

[7] Carlos P Roca, José A Cuesta, and Angel Sánchez. Effect of spatial structure on the evolution of cooperation. Physical Review E, 80(4):046106, 2009.

[8] Otso Ovaskainen, Dmitri Finkelshtein, Oleksandr Kutoviy, Stephen Cornell, Benjamin Bolker, and Yuri Kondratiev. A general mathematical framework for the analysis of spatiotemporal point processes. Theoretical ecology, 7(1):101–113, 2014.

[9] Rasmus Ibsen-Jensen, Krishnendu Chatterjee, and Martin A Nowak. Computational complexity of ecological and evolutionary spatial dynamics. Proceedings of the National Academy of Sciences, 112(51):15636–15641, 2015.

[10] Yu-Ting Chen. Wright–Fisher diffusions in stochastic spatial evolutionary games with death– birth updating. The Annals of Applied Probability, 28(6):3418–3490, 2018.

[11] Gisela García-Ramos and Diego Rodríguez. Evolutionary speed of species invasions. Evolution, 56(4):661–668, 2002.

[12] Vivian Hutson, Salomé Martinez, Konstantin Mischaikow, and Glenn T Vickers. The evolution of dispersal. Journal of mathematical biology, 47(6):483–517, 2003.

[13] Chris Cosner. Reaction-diffusion-advection models for the effects and evolution of dispersal. Discrete & Continuous Dynamical Systems, 34(5):1701, 2014.

[14] Bartlomiej Waclaw, Ivana Bozic, Meredith E Pittman, Ralph H Hruban, Bert Vogelstein, and Martin A Nowak. A spatial model predicts that dispersal and cell turnover limit intratumour heterogeneity. Nature, 525(7568):261–264, 2015.

[15] Li You, Joel S Brown, Frank Thuijsman, Jessica J Cunningham, Robert A Gatenby, Jingsong Zhang, andKateřina Staňková. Spatial vs. non-spatial eco-evolutionary dynamics in a tumor growth model. Journal of theoretical biology, 435:78–97, 2017.

[16] Jill A Gallaher, Pedro M Enriquez-Navas, Kimberly A Luddy, Robert A Gatenby, and Alexander RA Anderson. Spatial heterogeneity and evolutionary dynamics modulate time to recurrence in continuous and adaptive cancer therapies. Cancer research, 78(8):2127–2139, 2018.

[17] Diana Fusco, Matti Gralka, Jona Kayser, Alex Anderson, and Oskar Hallatschek. Excess of mutational jackpot events in expanding populations revealed by spatial Luria–Delbrück experiments. Nature communications, 7(1):1–9, 2016.

[18] Matti Gralka and Oskar Hallatschek. Environmental heterogeneity can tip the population genetics of range expansions. Elife, 8:e44359, 2019.

[19] Jakub Otwinowski and Joachim Krug. Clonal interference and Muller’s ratchet in spatial habitats. Physical biology, 11(5):056003, 2014.

[20] Matti Gralka, Fabian Stiewe, Fred Farrell, Wolfram Möbius, Bartlomiej Waclaw, and Oskar Hallatschek. Allele surfing promotes microbial adaptation from standing variation. Ecology letters, 19(8):889–898, 2016.

[21] Dominik Wodarz and Natalia L Komarova. Mutant evolution in spatially structured and fragmented expanding populations. Genetics, 216(1):191–203, 2020.

[22] Hisashi Ohtsuki and Martin A Nowak. The replicator equation on graphs. Journal of theoretical biology, 243(1):86–97, 2006.

[23] Corina E Tarnita, Hisashi Ohtsuki, Tibor Antal, Feng Fu, and Martin A Nowak. Strategy selection in structured populations. Journal of theoretical biology, 259(3):570–581, 2009.

[24] Corina E Tarnita, Nicholas Wage, and Martin A Nowak. Multiple strategies in structured populations. Proceedings of the National Academy of Sciences, 108(6):2334–2337, 2011.

[25] Rick Durrett. Spatial evolutionary games with small selection coefficients. Electronic Journal of Probability, 19:1–64, 2014.

[26] Wolfgang Stephan. Selective sweeps. Genetics, 211(1):5–13, 2019.

[27] Priyanka Gopal, Elif Irem Sarihan, Eui Kyu Chie, Gwendolyn Kuzmishin, Semihcan Doken, Nathan A Pennell, Daniel P Raymond, Sudish C Murthy, Usman Ahmad, Siva Raja, et al. Clonal selection confers distinct evolutionary trajectories in BRAF-driven cancers. Nature communications, 10(1):1–14, 2019.

[28] Frederick M Cohan. Bacterial speciation: genetic sweeps in bacterial species. Current Biology, 26(3):R112–R115, 2016.

[29] Lauren MF Merlo, John W Pepper, Brian J Reid, and Carlo C Maley. Cancer as an evolutionary and ecological process. Nature reviews cancer, 6(12):924–935, 2006.

[30] Daniel A Haber and Jeff Settleman. Drivers and passengers. Nature, 446(7132):145–146, 2007.

[31] Mel Greaves and Carlo C Maley. Clonal evolution in cancer. Nature, 481(7381):306–313, 2012.

[32] Matthew H Bailey, Collin Tokheim, Eduard Porta-Pardo, Sohini Sengupta, Denis Bertrand, Amila Weerasinghe, Antonio Colaprico, Michael C Wendl, Jaegil Kim, Brendan Reardon, et al. Comprehensive characterization of cancer driver genes and mutations. Cell, 173(2):371–385, 2018.

[33] Natalia L Komarova, Jan A Burger, and Dominik Wodarz. Evolution of ibrutinib resistance in chronic lymphocytic leukemia (CLL). Proceedings of the National Academy of Sciences, 111 (38):13906–13911, 2014.

[34] Dominik Wodarz, Naveen Garg, Natalia L Komarova, Ohad Benjamini, Michael J Keating, William G Wierda, Hagop Kantarjian, Danelle James, Susan O’ Brien, and Jan A Burger. Kinetics of CLL cells in tissues and blood during therapy with the BTK inhibitor ibrutinib. Blood, The Journal of the American Society of Hematology, 123(26):4132–4135, 2014.

